# Efference copy is necessary for the attenuation of self-generated touch

**DOI:** 10.1101/823575

**Authors:** Konstantina Kilteni, Patrick Engeler, H. Henrik Ehrsson

**Affiliations:** Department of Neuroscience, Karolinska Institutet, Solnavägen 9, 17165 Stockholm, Sweden

**Keywords:** Somatosensory attenuation, efference copy, passive movements, internal models, sensorimotor predictions

## Abstract

A self-generated touch feels less intense than an external touch of the exact same intensity. According to a prevalent computational theory of motor control, this attenuation occurs because the brain uses internal forward models to predict the somatosensory consequences of our movements using a copy of the motor command, i.e., the efference copy. These tactile predictions are then used to suppress the perceived intensity of the actual tactile feedback. Despite being highly influential, the core assumption of theory has never been tested; that is, whether the efference copy is necessary for somatosensory attenuation. A possible alternative hypothesis is that a predictable contact of two of one’s own body parts is sufficient. Using a psychophysical task, we quantified the attenuation of touch applied on the participants’ left index finger when the touch was triggered by the active or passive movement of the participants’ right index finger and when it was externally generated in the absence of any movement. We observed somatosensory attenuation only when the touch was triggered by the voluntary movement of the participants’ finger. In contrast, during the passive movement, the intensity of the touch was perceived to be as strong as when the touch was externally triggered. In both active and passive movement conditions, the participants showed the same discrimination capacity. Electromyographic recordings confirmed minimal activity of the right hand during the passive movement. Together, our results suggest that the efference copy is necessary for computing the somatosensory predictions that produce the attenuation of self-generated touch.

## Introduction

Somatosensory attenuation refers to the phenomenon wherein a self-generated touch feels weaker than an externally generated touch of the same intensity. Several behavioral experiments have shown that participants judge a tap or a stoke delivered on their relaxed hand as less intense when the touch is produced by the active movement of their other hand compared to when it is produced externally by a motor (Bays *et al.*, 2005; Blakemore *et al.*, 1999; Kilteni *et al.*, 2019). Similarly, when participants were asked to match external forces applied to their relaxed index fingers by reproducing the same forces with their other index fingers through bimanual action simulating direct contact between the digits (force-matching task), they produced stronger forces than the ones required; this is because the self-generated forces are being perceptually attenuated (Kilteni *et al.*, 2018; Kilteni and Ehrsson, 2017b; a; Shergill *et al.*, 2003).

Motor control theories suggest that somatosensory attenuation arises from the same predictive processes that the brain uses when planning and executing movements, the so-called internal models. Accordingly, when we perform a movement, the internal model uses a copy of the motor command (i.e., the efference copy) to predict the sensory (including the somatosensory) consequences of our movements. These predictions are then used to compensate for the intrinsic delays in receiving sensory feedback (Davidson and Wolpert, 2005; Franklin and Wolpert, 2011; Kawato, 1999) but also to attenuate the self-generated somatosensory signals and thus to increase the salience of any externally generated tactile information (Bays and Wolpert, 2007; Blakemore *et al.*, 2000). The internal models have been suggested to be located in the cerebellum (Shadmehr *et al.*, 2010; Shadmehr and Krakauer, 2008; Therrien and Bastian, 2018; Wolpert *et al.*, 1998), and neuroimaging studies on somatosensory attenuation have indeed revealed cerebellar activity when comparing conditions that include externally generated touches with those that include self-generated touches (Blakemore et al. 1998; Kilteni and Ehrsson *Under Review*).

The importance of the efference copy for somatosensory attenuation is well established within the motor control community. Indeed, all previously mentioned behavioral studies of somatosensory attenuation (Bays *et al.*, 2006, 2005; Bays and Wolpert, 2008; Kilteni *et al.*, 2018, 2019; Kilteni and Ehrsson, 2019, 2017b; a; Palmer *et al.*, 2016; Shergill *et al.*, 2005, 2014, 2003; Walsh *et al.*, 2011; Wolpe *et al.*, 2016) use conditions with voluntary movement, and it is generally assumed that it is the efference copy associated with the voluntary motor commands that is critical for the attenuation phenomenon to occur. However, this assumption has not been directly tested. This is problematic because the experimental conditions that produce somatosensory attenuation not only involve efference copy but also the *prediction* and the *perception* of self-touch. For example, in the classic force-matching task, when participants press one index finger against the other and somatosensory attenuation is observed, this includes the efference copy, the prediction of contact between the hands and the perceptual experience from the bimanual interaction. Thus, a parsimonious alternative model for somatosensory attenuation is that the mere prediction and perception of self-touch between two of one’s own body parts could be the critical factor that triggers the phenomenon and not the efference copy.

To the best of our knowledge, the results of all previously published studies on sensory attenuation using bimanual force-matching tasks would be consistent with this alternative view. In line with this, if a distance is introduced between the two fingers that makes both unlikely and non-feasible the physical contact of the digits in the force-matching task, the attenuation is eliminated or significantly reduced (Bays and Wolpert, 2008; Kilteni and Ehrsson, 2017b). Moreover, it is the *prediction* and *perception* of self-touch that is important, not the actual contact between the hands; this was demonstrated in experiments where the participants experienced the illusion where a plastic right hand seen to press against their left hand was thought to be their own right hand (rubber hand illusion), which led to an attenuation of the forces even though their real hand was kept at a distance from the right hand (Kilteni and Ehrsson, 2017a). Furthermore, the stronger the illusion that the participants experienced was, the stronger the attenuation of the self-produced forces. Further support on the importance of the prediction of self-touch can be found in the study of Bays et al. (2006) who observed somatosensory attenuation also when the participants’ hands unexpectedly failed to touch each other. All these findings have previously been interpreted in a theoretical model in which the internal model uses both the efference copy and information about the sensory state of the body to compute the likelihood of self-touch and the associated attenuation (Blakemore *et al.*, 2000; Kilteni and Ehrsson, 2017a) (Figure 1). According to the alternative theory, however, the brain would attenuate self-touch through sensory predictions that are purely based on (i) the sensory state of the body, indicating that one hand is (likely) directly touching the other hand, and (ii) the belief that the touch is caused by this single event of the two own body parts contacting each other (Figure 1). This generalized predictive mechanism does not consider the efference copy as a prerequisite, and it relates to the predictive coding theory that state that the brain forms predictions based on its prior beliefs and continuously updates them to minimize any error between the predicted and the incoming sensory information (Friston, 2005, 2009; Rao and Ballard, 1999). Moreover, this theory is supported by earlier observations that neural responses become suppressed after the repeated presentation of a stimulus (repetition suppression) or after the presentation of an expected stimulus (for a review see (Grotheer and Kovács, 2016)). Importantly, this theory would not necessarily speak against the internal models’ theory, but it would favor a universal predictive mechanism underlying all multisensory bodily events, including somatosensory attenuation that is not necessarily based on motor signals; the predictions of this mechanism could be more finely tuned when a motor command is available.

**Figure 1.**
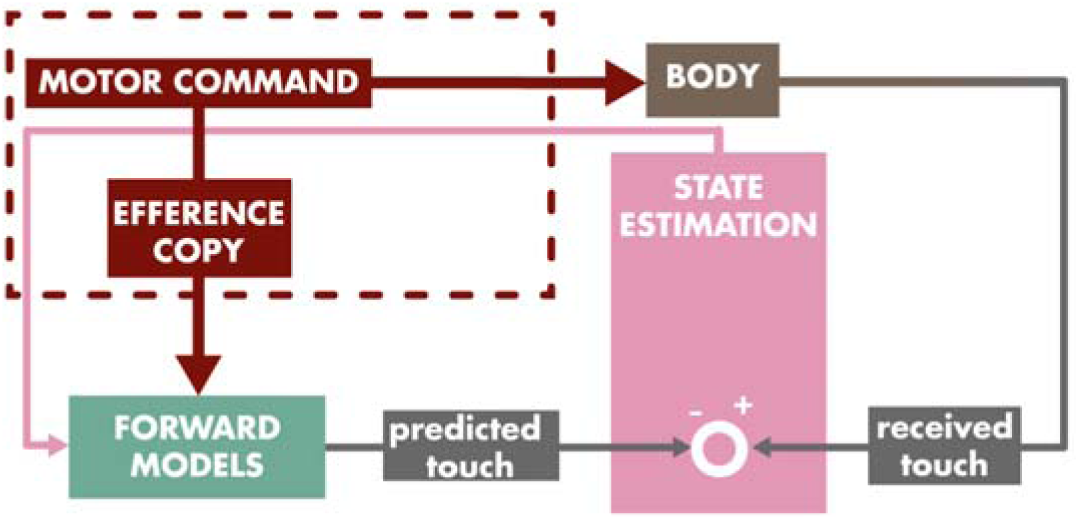
A theoretical model for somatosensory attenuation. According to the efference copy-based theoretical model, during the active movement of the right hand to touch the left hand, a copy of the motor command discharged to the right hand (the efference copy) is sent to the forward model that predicts the next state (e.g., position) of the right hand as well as the sensory consequences associated with that state (e.g., proprioceptive input). Similarly, the next state of the left hand is predicted, although this should remain motionless. Predicted and incoming information are combined in the state estimation process. If the predicted positions of the two hands are close, touch is additionally predicted and thus the incoming touch is attenuated. According to the alternative hypothesis describing a general predictive mechanism underlying somatosensory attenuation in the absence of the efference copy, during the passive movement of the right hand towards the left hand, there is no motor command and, thus, no efference copy (dark red part is absent from the model). The incoming sensory input (e.g., proprioception) is used in combination with prior beliefs from the forward models (“where I expect my hand to be”) to estimate the states of the two hands. The estimated states are fed back to the forward models. As before, if the predicted states of the two hands are close, touch is predicted and the incoming touch becomes attenuated. The present study investigated whether the motor command and thus the efference copy (the part of model denoted by the dark red dotted line) is a prerequisite of this predictive attenuation mechanism to dissociate between these two models.

Here, we used a psychophysics paradigm (Bays *et al.*, 2005; Kilteni *et al.*, 2019) to quantitatively compare somatosensory attenuation in conditions with active and passive movements to directly test the hypothesized necessary role of the efference copy in the attenuation of self-touch and thus to distinguish between the two alternative hypotheses discussed above. The passive movement of one index finger to touch the other lacks the efference copy but does involve the prediction and perception of self-touch. Therefore, if somatosensory attenuation is observed only when the touch is produced by a voluntary movement (active movement), this would indicate that the efference copy is necessary and it would speak in favor of the internal models’ theory. Alternatively, if somatosensory attenuation is also observed during a passive movement, this would support the generic multisensory predictive model of attenuation.

## Materials and Methods

### Participants

After providing written informed consent, thirty participants (15 women and 15 men, 29 right-handed and 1 left-handed) aged 18-39 years participated in the present study. The sample size was decided based on a previous study using the same task (Kilteni *et al.*, 2019). Handedness was assessed using the Edinburgh Handedness Inventory (Oldfield, 1971). The Swedish Ethical Review Authority (https://etikprovningsmyndigheten.se/) approved the study (no. 2016/445-31/2, amendments 2018/254-32 and 2019-03063).

### Materials and Procedures

Participants were asked to place their left index finger inside a molded support while their right arm comfortably rested on top of a set of sponges. In each trial, a DC electric motor (Maxon EC Motor EC 90 flat; manufactured in Switzerland) delivered two taps (the test tap and the comparison tap in Figure 2a-c) on the pulp of the participants’ left index finger through a cylindrical probe (25□mm height) with a flat aluminum surface (20□mm diameter) attached to the motor’s lever. A force sensor (FSG15N1A, Honeywell Inc.; diameter, 5 mm; minimum resolution, 0.01 N; response time, 1 ms; measurement range, 0–15 N) was placed within the probe to record the forces applied on the left index finger (red sensor in Figure 2a-c).

**Figure 2.**
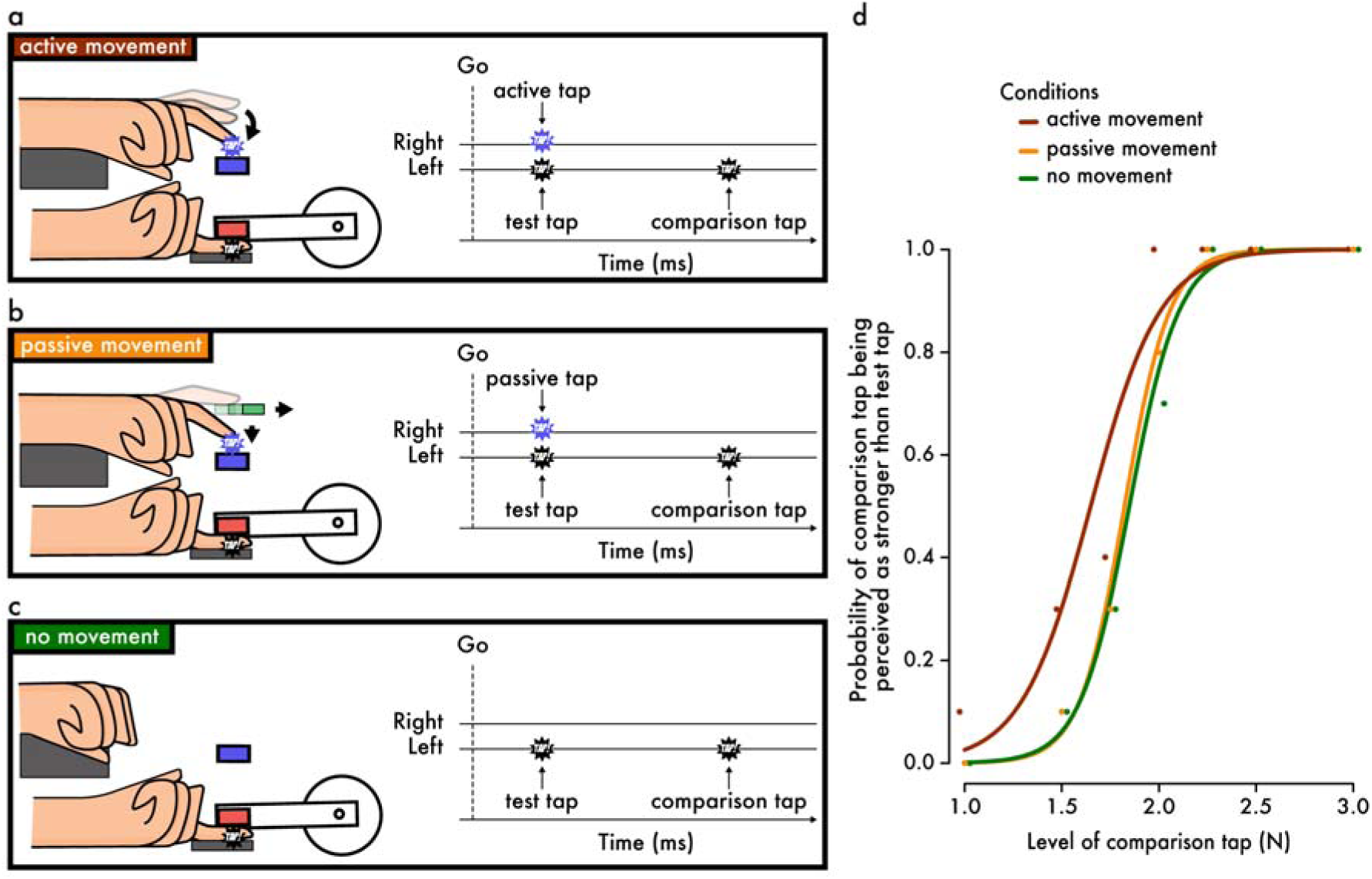
Experimental setup, design and analysis. In all three conditions, the participants received two taps on their relaxed left index finger (a test tap and a comparison tap), and they were requested to indicate which tap felt stronger. In the active movement condition (**a**), the participants actively tapped a force sensor with their right index finger (blue sensor). This active tap simultaneously triggered the test tap on the participants’ left index finger. In the passive movement condition (**b**), the participants’ right index finger was left to fall on the force sensor (blue sensor) and passively tap it. This passive tap simultaneously triggered the test tap on the participants’ left index finger. In the no movement condition (**c**), the participants remained relaxed, and the test tap was externally triggered. (**d**) Data from a representative participant. For each condition, the participant’s responses were fitted with psychometric curves, and the point of subjective equality (PSE) and the just noticeable difference (JND) were extracted. We have horizontally jittered the points to avoid their overlapping.

In the *active movement* condition (Figure 2a), participants were asked to actively tap with their right index finger a force sensor (same specifications as above) placed on top of (but not in contact with) the probe upon an auditory ‘go’ cue (blue sensor in Figure 2a-c). Participants were asked to tap the sensor after the ‘go’ cue, neither too hard nor too softly but “as strongly as when they tap the surface of their smartphone”. Their active tap on the force sensor triggered the test tap with an intrinsic delay of 36 ms (threshold set to 0.15 N).

In the *passive movement* condition (Figure 2b), participants were asked to rest their right index finger on top of a plastic surface that was placed on top of (but not in contact with) the sensor for the right index finger. Upon an auditory ‘go’ cue, a servomotor (Hitec HS-81) retracted this surface away, and the participants’ right index finger freely fell on the underlying sensor. As before, the passive tap on the force sensor (> 0.15 N) triggered the test tap with a minimal (36 ms) delay. Significant training took place before this condition to ensure that the participants did not resist the action and did not produce any large muscular activity, as well as to confirm that the finger fell freely on the sensor. To minimize the elicitation of any motor reflexes due to surprise, the passive movement condition was designed to be as predictable as possible by retracting the surface always at the same time after the ‘go’ cue.

In the *no movement* condition (Figure 2c), participants kept their right hand on top of the sponges. After the auditory ‘go’ cue, the test tap was applied to the participants’ left index finger.

In all conditions, the view of the pulp of the left index finger was occluded, and participants were asked to fixate on a cross placed on a wall 2 m opposite them. A force of 0.1 N was constantly applied on the participants’ left index finger to avoid overshooting in the experimental forces. Any sounds created by the motor, by the right hand’s tap, or by the servomotor were precluded by administering white noise to the participants through a pair of headphones. No feedback was provided to the participants. EMG was recorded from the right first dorsal interosseous muscle (FDI) (see below for details). The order of conditions was fully counterbalanced across participants. The experiment lasted 60 minutes approximately.

After the end of the three conditions, all participants were asked whether they spontaneously performed motor imagery during the passive movement condition. We asked this question to exclude the putative concern that participants would spontaneously engage in mental simulation in this condition, which would produce somatosensory attenuation through an efference copy-based mechanism (Kilteni *et al.*, 2018).

### Psychophysics

Each condition involved 70 trials. The test tap was set to 2 N, while the intensity of the comparison tap was systematically varied among seven different force levels (1, 1.5, 1.75, 2, 2.25, 2.5 or 3 N). The two taps had a 100 ms duration, and the delay between them was random (800 – 1500 ms). On every trial, participants had to verbally indicate which tap on their left index finger felt stronger: the first (test) or the second (comparison). They were told that they should not try to balance their responses (50% first and 50% second) and they were asked to guess when the intensity of the two taps felt very similar.

For each condition, the participants’ responses were fitted with a generalized linear model using a *logit* link function (Equation 1, Figure 2d).

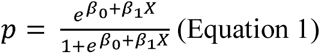

Two parameters of interest were extracted: the point of subjective equality 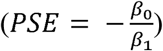, which represents the intensity at which the test tap felt as strong as the comparison tap (*p* = 0.5) and which quantifies the attenuation, and the just noticeable difference 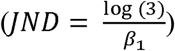, which reflects the participants’ sensitivity for the force discrimination.

During the data collection, trials during which the right index finger was seen not to fall properly were rejected and repeated to reach 70 trials per condition. After the data collection, twenty-six force trials (26 of 6300, 0.4%) were rejected: in five trials, the responses were missing; in three trials, the intensity of the test tap (2 N) was not applied correctly; and in eighteen, the passive movement was not properly performed as instructed. These 26 trials were also rejected from the EMG. Before fitting the responses, the values of the applied comparison taps were binned to the closest value with respect to their theoretical values (1, 1.5, 1.75, 2, 2.25, 2.5 or 3 N).

### EMG acquisition and preprocessing

Surface EMG was recorded using the Delsys Bagnoli electromyography system (DE-2.1 Single Differential Electrodes) from the belly of the right FDI muscle after cleaning the skin with alcohol. The EMG reference electrode was placed either on the left clavicle or on the superior anterior iliac spine. The signals were analog bandpass filtered between 20 and 450☐Hz, sampled at 2.0☐kHz and amplified (gain ☐=☐1000). EMG data were preprocessed in MATLAB. A bandstop filter was used to suppress the 50☐Hz powerline interference, and the DC offsets of the signals were removed.

### EMG analysis

For each trial, we calculated the root mean square (RMS) of the EMG signal during the time window from the ‘go’ cue to the test tap. The window length in the *active movement* condition depended on the participants’ reaction time to tap the sensor and was 716.8 ± 186.8 ms (mean ± sd). For the *passive movement* condition, the duration of the windows could slightly change depending on how the participants placed the finger on the surface and was 287.1 ± 36.7 ms. Finally, in the *no movement* condition, the duration of the windows was fixed at 599.8 ± 0.2 ms. We averaged the RMS activity across all (70) trials and then compared the mean RMS across participants between the three conditions.

During data collection, trials in which the participants did not relax their right index finger (in the *passive* and *no movement* conditions) or where there was visibly larger EMG activity during the test tap compared to the comparison tap (for the *passive* condition) were rejected and repeated. For one participant, the EMG data from the *active movement* condition were not registered; thus, the EMG analysis was performed with 29 subjects.

### Statistical analysis

Data were analyzed using R (Core Team, 2018) and JASP (JASP Team 2019). The normality of the PSE, the JND and the EMG data distributions was checked using the Shapiro-Wilk test. Depending on their normality, we performed planned comparisons using either a paired t-test or a Wilcoxon signed-rank test. We report 95% confidence intervals (*CI^95^)* for each statistical test. Effect sizes are given by Cohen’s *d* if the data were normally distributed or by the matched rank biserial correlation *r*_*rb*_ if the data were not normally distributed. In addition, a Bayesian factor analysis using default Cauchy priors with a scale of 0.707 was carried out to provide information about the level of support for the alternative hypothesis compared to the null hypothesis (*BF_10_*) given the data. Finally, a correlation was tested using Kendall’s Tau-b coefficient τ_*B*_ given that the data were not normally distributed. All tests were two-tailed.

## Results

Figure 3 shows the average and individual PSEs extracted for each condition, as well as the individual differences per pair of conditions. In agreement with previous studies (Bays *et al.*, 2005; Kilteni *et al.*, 2019), a tap that was self-generated through a voluntary movement (*active movement* condition) felt significantly weaker compared to an externally generated identical tap (*no movement* condition): Wilcoxon signed rank test, *n* = 30, *V* = 6, *p* < 0.001, *CI*^95^ = [-0.268, −0.156], *r*_*rb*_ = −0.974, *BF*_10_ > 14246. This is the classic phenomenon of somatosensory attenuation. Importantly, the self-generated tap (*active movement* condition) was significantly attenuated compared to the tap of the same intensity that resulted from passive movement (*passive movement* condition): Wilcoxon signed rank test, *n* = 30, *V* = 8, *p* < 0.001, *CI*^95^ = [-0.286, −0.144], *r*_*rb*_ = −0.966, *BF*_10_ > 1325. Notably, the perception of the tap that resulted from passive movement (*passive movement* condition) did not significantly differ from that of the externally generated tap (*no movement* condition), and the Bayesian analysis indicated that the level of perceived force was similar in the two conditions: paired t-test, *n* = 30, *t*(29) = 0.26, *p* = 0.799, *CI*^95^ = [-0.064, 0.083], Cohen’s *d* = 0.047, *BF*_10_ = 0.20. Collectively, these results suggest that only the somatosensory feedback from the self-generated taps is attenuated.

**Figure 3.**
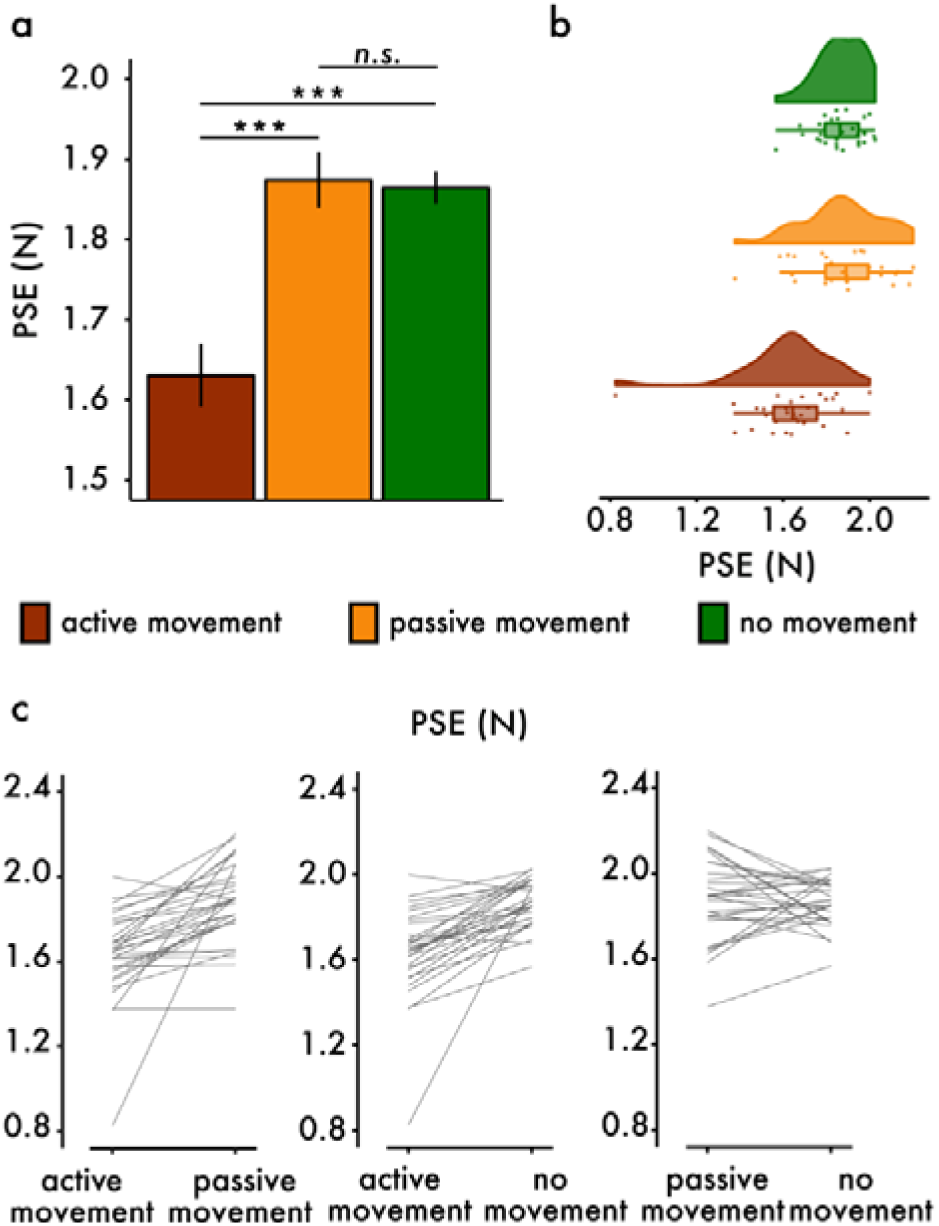
Results on the points of subjective equality (PSEs). (**a**) Bar graphs show the PSEs (mean ± se) per condition (*\*\*\* p* < 0.001, *n.s.* not significant). Only the *active movement* condition produced somatosensory attenuation. In contrast, no changes were detected in the PSEs between the *passive movement* and the *no movement* condition. (**b**) Raincloud plots (Allen *et al.*, 2019) show the raw PSEs as well as their distribution per condition. (**c**) Line plots illustrate the participants’ paired responses per combination of conditions.

Figure 4 shows the average and individual JNDs extracted for each condition, as well as the individual differences per pair of conditions. Participants showed similar response sensitivities in the force discrimination task between the *active movement* and the *passive movement* conditions, ruling out the possibility that one condition was more or less difficult than the other: paired t-test, *n* = 30, *t*(29) = 0.42, *p* = 0.680, *CI*^95^ =[-0.024, 0.036], Cohen’s *d* = 0.076, *BF*_10_ = 0.211. Both the *active movement* (paired t-test, *n* = 30, *t*(29) = 2.25, *p* = 0.032, *CI*^95^ =[0.003, 0.065], Cohen’s *d* = 0.411, *BF*_10_ = 1.706) and *passive movement* conditions (paired t-test, *n* = 30, *t*(29) = 2.11, *p* = 0.044, *CI*^95^ =[0.001, 0.055], Cohen’s *d* = 0.384, *BF*_10_ = 1.323) showed significantly lower discrimination capacities than the *no movement* condition. The Bayesian analysis did not provide any conclusive support for the existence of such differences (*BF*_10_ < 2 in both cases) and thus, one should be cautious on interpreting the frequentist analysis. Nevertheless, if these JND differences do exist, they indicate that the movement of the right index finger per se, either voluntary or not, deteriorates the discrimination performance on the left index finger. This because in both *active* and *passive* movement conditions, the participants had to direct their attention to both hands (i.e., the movement of the right index and the force discrimination task on the left index), while in the *no movement* condition, the participants directed their attention only to the left index finger. Another related factor could be the presence of sensory feedback on the right index finger simultaneous to the sensory feedback on the left hand in the movement conditions that could render the task slightly more demanding.

**Figure 4.**
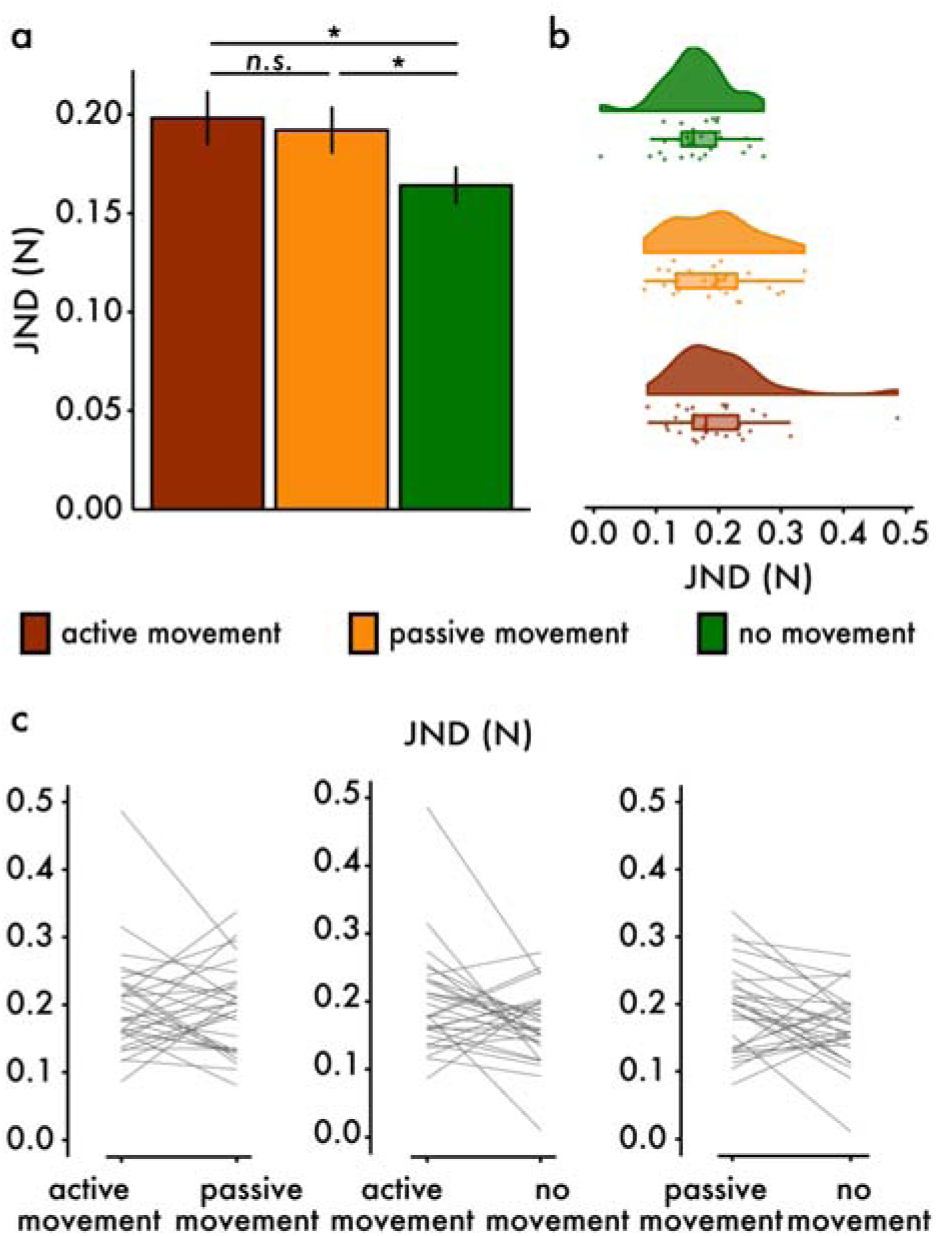
Results on the just noticeable difference (JNDs). (**a**) Bar graphs show the JNDs (mean ± se) per condition (*\*p* < 0.05, *n.s.* not significant). The *active* and *passive movement* conditions showed higher JND than the *no movement* condition. In contrast, no changes were detected in the JNDs between the *active* and *passive movement* condition. (**b**) Raincloud plots (Allen *et al.*, 2019) show the raw JNDs as well as their distribution per condition. (**c**) Line plots illustrate the participants’ paired responses per combination of conditions.

Figure 5 shows the group psychometric functions per condition using the corresponding mean PSE and JND (see also Appendix Supplementary Figure S1 for all individual fits). Somatosensory attenuation was produced only during a self-generated (voluntary) movement.

**Figure 5.**
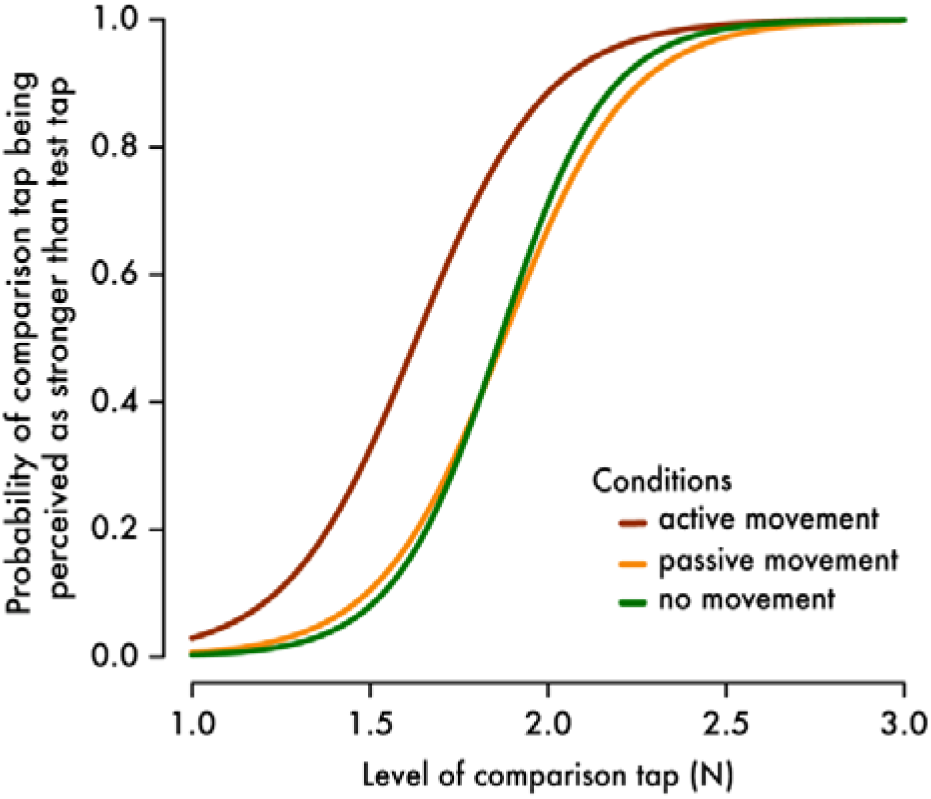
Group psychometric functions per condition. The plots were generated using the mean PSE and the mean JND across the thirty participants per condition. Significant attenuation with respect to the *no movement* condition was observed only in the *active movement* condition.

It should be noted that the *active* and *passive movement* conditions differed not only in terms of the efferent signals discharged to the right index finger for pressing but also in terms of the afferent somatosensory feedback received from the right index finger; that is, the force that was applied by the sensor to the right index finger, opposite to the pressing force. The participants pressed stronger forces with their right index finger during the *active* (*mean* ± *sd*: 1.210 ± 0.790 N) than during the *passive movement* condition (0.431 ± 0.134 N): Wilcoxon signed rank test, *n* = 30, *V* = 455, *p* < 0.001, *CI*^95^ = [0.441, 0.988], *r*_*rb*_ = 0.957, *BF*_10_ > 3145. To rule out the unlikely possibility that passive movements did not produce somatosensory attenuation because of the reduced force and somatosensory feedback from the right index finger, we tested for a relationship between the forces the participants pressed on the sensor (passive tap, Figure 2) and their PSEs in the passive condition. As we expected, no relationship was found: *n* = 30, *T* = 205, τ_*B*_ = −0.057, *p* = 0.671, *CI*^95^ = [-0.279, 0.164], with the Bayesian analysis favoring the null hypothesis: *BF*_10_ = 0.259. We further performed the same analysis with the JNDs; no relationship was found between the JND in the passive movement condition and the somatosensory feedback from the right index finger: *n* = 30, *T* = 235, τ_*B*_ = 0.080, *p* = 0.547, *CI*^95^ = [-0.201, 0.362], with the Bayesian analysis favoring again the null hypothesis: *BF*_10_ = 0.284.

Next, we analyzed the EMG data to test whether participants were relaxed during the passive movement condition. Figure 6a and b shows the average and individual RMS activity calculated per condition, and Figure 6c illustrates the individual differences per pair of conditions. Validating our experimental manipulation, analysis of the RMS activity revealed significantly higher activity in the *active movement* condition compared to the *no movement* condition (Wilcoxon signed rank test, *n* = 29, *V* = 435, *p* < 0.001, *CI*^95^ =[0.048, 0.084], *r*_*rb*_ = 1, *BF*_10_ > 4.48 × 10^6^) and the *passive movement* condition (Wilcoxon signed rank test, *n* = 29, *V* = 435, *p* < 0.001, *CI*^95^ =[0.046, 0.084], *r*_*rb*_ = 1, *BF*_10_ > 3.27 × 10^6^). The *passive movement* condition did reveal small EMG activity compared to the *no movement* condition (Wilcoxon signed rank test, *n* = 29, *V* = 394, *p* < 0.001, *CI*^95^ = [0.0003, 0.001], *r*_*rb*_ = 0.811, *BF*_10_ = 8.257), but this increase was ≅ 70 times smaller compared to the increase in the *active movement* condition (Figure 6c). Thus, we conclude that the participants were able to relax in the passive condition and that the experimental comparison of active versus passive finger movements was successfully implemented in our paradigm.

**Figure 6.**
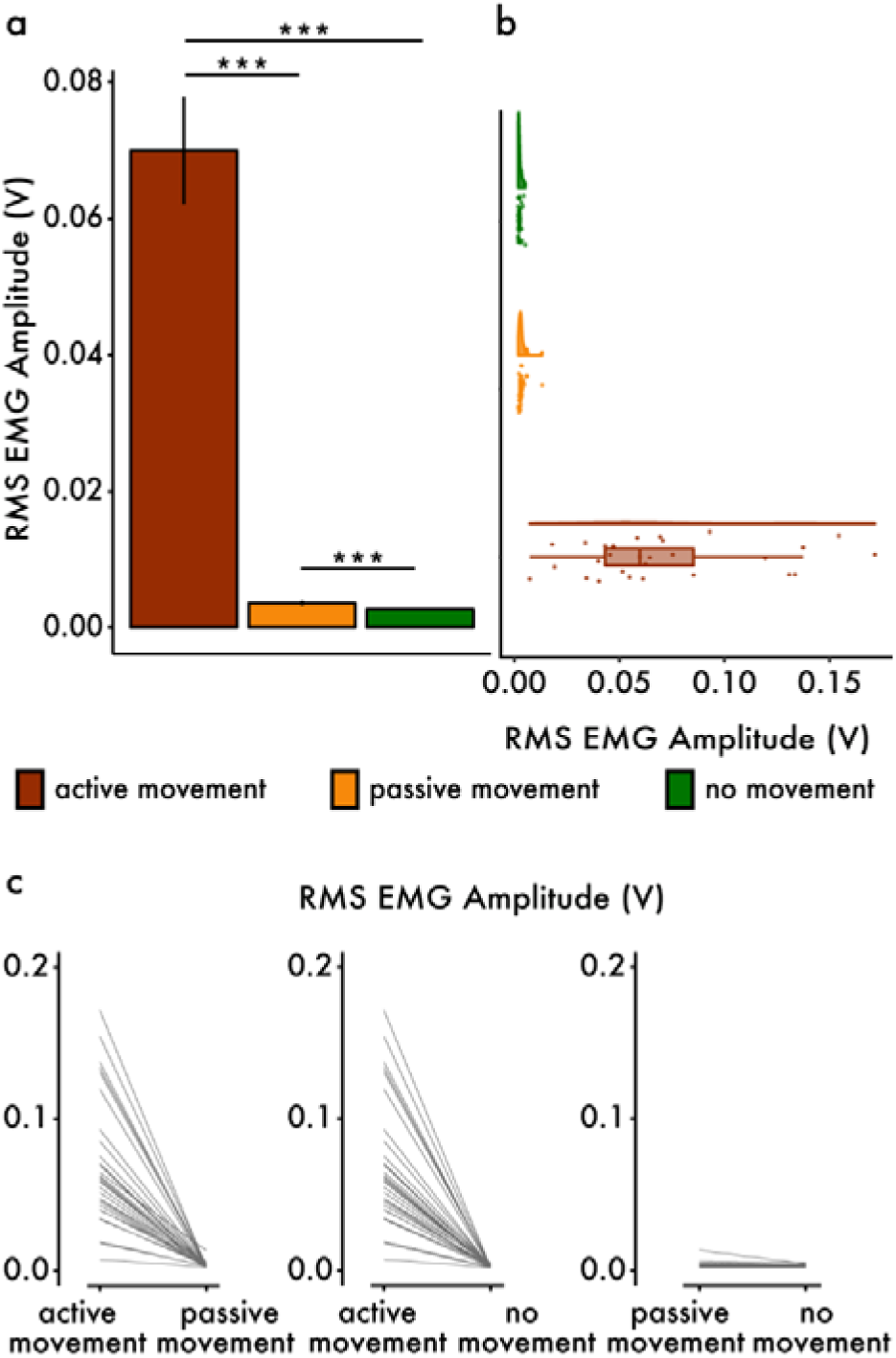
Results on the EMG RMS amplitude. (**a**) Bar graphs show the mean RMS amplitude (± SE) per condition (*\*\*\* p* < 0.001). (**b**) Raincloud plots show the raw amplitudes as well as their distributions per condition. (**c**) Line plots show the participants’ paired responses per combination of conditions.

Finally, with respect to the motor imagery question, none of the thirty participants reported performing motor imagery during the passive movement condition. This excludes the possibility that the *passive movement* condition was confounded with motor simulation and thus with efference copies – a factor that could drive somatosensory attenuation *per se* (Kilteni *et al.*, 2018).

## Discussion

The present study found that touch applied on a static left index finger gets attenuated only when it results from the active movement of the right index finger, not when it results from the passive movement of the right index finger or when it is applied in the absence of any movement. Specifically, the perceived intensity of a touch that results from the passive movement of the right index finger was comparable to that of an externally generated touch. These findings favor the interpretation based on the internal models and suggest that the efference copy is necessary for the attenuation of self-generated touch. According to this theory, during the *active movement* condition, a copy of the motor command sent to the right hand (the efference copy) is used to predict the next state (e.g., position) of the hand and its expected sensory consequences associated with that state (Bays and Wolpert, 2008; Blakemore *et al.*, 2000; Wolpert and Flanagan, 2001; Wolpert and Ghahramani, 2000). Since the predicted end-position of the right index finger falls very close to that of the relaxed left index finger, touch is predicted on this left finger as well (Kilteni and Ehrsson, 2017b). The actual touch (here, the test tap) is attenuated once it is received since it has been predicted based on the efference copy from the motor command to the right index finger. From a computational perspective, the present study demonstrates that it is the voluntary direct contact of the two body parts that is critical for somatosensory attenuation and not the mere contact or close proximity between the two involved body parts produced by the (active/passive) movement. This supports the internal model theory of sensory attenuation and speaks against the general multisensory predictive hypothesis.

We first discuss three methodical issues: (i) were the active and passive tasks comparable in terms of performance on the discrimination task and the predictability of touch? (ii) was the passive task free of efference copies? and (iii) could small differences in tactile feedback from the right index finger between the active and the passive movement conditions influence the somatosensory attenuation on the left index? With respect to the first question, it is important to stress that there were no task differences between the *active* and *passive movement* conditions that could influence the participants’ responses in the force discrimination task. First, the two conditions had similar JNDs, suggesting that the participants’ performance sensitivity did not differ between the two conditions (Figures 3 and 4). Second, we designed the *passive movement* condition to minimize any surprises and make it as predictable as possible, similar to the *active movement* condition. Specifically, in the *passive movement* condition, the platform was always retracted at the same time to facilitate the anticipation of the timing of the hands’ contact and to strengthen the causal link between the passive displacement of one finger and the somatosensory input of the other finger – as in the *active movement* condition. With respect to the sensory predictability, an earlier study on the unloading task (Diedrichsen *et al.*, 2003) showed that anticipatory adjustments are present only when the efference copy is available; in contrast, no adjustments were observed in the absence of a voluntary movement, even when the predictability of the sensory stimulus was high. Therefore, in the present study the absence of attenuation in the passive movement condition suggests that the motor system cannot predict the consequences of an involuntary movement as precisely as those of a voluntary one because of the lack of efference copy.

With respect to the second question, it is noteworthy that the *passive movement* condition did yield some muscular activity compared to the *no movement* condition, but its magnitude was much (approximately 70 times) smaller than the one elicited in the *active movement* condition. This weak muscular activity in the passive condition could represent reflexes for automatic postural stabilization or stretch reflexes (Doemges and Rack, 1992) rather than voluntary motor commands. Importantly, this interpretation is in line with the fact that we did not observe any reliable somatosensory attenuation in the passive condition. Another related putative concern is that the participants might spontaneously start to imagine active movements in the passive condition. We know that imagery of voluntary self-touch can lead to somatosensory attenuation, presumably by engaging the efference copy when internally simulating the action (Kilteni et al. 2018). As an extra precaution to rule out this unlikely scenario, we asked our participants to indicate whether they performed motor imagery during the passive movement, and they all denied doing so. Therefore, we can exclude the possibility that participants mentally simulated an active movement in the passive condition (Kilteni *et al.*, 2018). Thus, we think it is reasonable to conclude that the passive condition was free of efferent copies, at least to the extent that matters for the interpretation of the results.

The third concern was that the *passive movement* condition also differed from the *active movement* condition in terms of the somatosensory feedback received from the *right* index finger because the subjects pressed smaller forces with their right index finger in the *passive* compared to the *active movement* condition. We did not find any evidence that this reduced feedback could hinder somatosensory attenuation during passive movements. Further evidence comes from a previous somatosensory attenuation study that used the same psychophysics task as the present study; in the study of Bays et al. (2005) participants did not move their right index finger but they received a tap from an upward force pulse at the same time they received the tap on their left index finger that was of similar magnitude. Despite the enhanced somatosensory feedback, the participants did not show any attenuation. Moreover, an earlier study on the force-matching task found no effect on somatosensory attenuation by different relationships (gains) between the forces participants pressed with their right index finger and the forces they received on their left index finger, as long as this relationship were stable (Bays and Wolpert, 2008). This further corroborates the hypothesis that the somatosensory feedback from the right index finger *per se* is not critical for somatosensory attenuation on the left index finger in the bimanual force matching task.

It is interesting to consider the present results together with the findings that were recently reported by Kilteni et al. (2018) on somatosensory attenuation during motor imagery. Motor imagery corresponds to internally simulating movement without executing it, which involves producing a central motor command and thus efference copy. In that study, Kilteni et al. (2018) asked their participants to imagine pressing their right index finger against their left index finger through a sensor while they simultaneously received a force on their left index finger. The experimenters observed that when the tactile consequences of the imagined movement matched the received touch in terms of space and time, the touch was attenuated to the same extent as when the participants actually executed the movement. This result suggests that the efference copy is sufficient for somatosensory attenuation when the sensory predictions derived from the efference copy are spatiotemporally congruent with the actual somatosensory feedback. Importantly, the present results add to this by suggesting that the efference copy is not only sufficient but also *necessary* for sensory attenuation of self-touch, which has important bearings on the computational models of sensory attenuation.

The difference between active and passive movements in terms of perceptual stability and sensory processing has been shown in modalities other than touch. For example, in their seminal observations within the visual domain, first Descartes and later Helmholtz (1867) noted that when we actively move our eyes, the world seems stable; in contrast, when we tap the side of our eyeball to ‘passively’ change the retinal image, the world appears to be moving. That is, the visual consequences of this passive displacement are processed differently from those produced by active eye movements. Similarly, in primates, it has been systematically shown that the vestibular consequences of active head movements are significantly attenuated, in striking contrast to vestibular information received during passive head movements (see (Cullen, 2012) for a review). Our findings provide evidence that a similar distinction applies in the somatosensory domain as well: only touches that result from an active and not from a passive movement become attenuated.

Efference copy-related circuits have been revealed in several species across the animal kingdom (Crapse and Sommer, 2008), suggesting that predictive signals computed from the motor command might constitute a generalized strategy for biological organisms to differentiate self-generated from externally generated information. The present study showed that only active movements allow the computations of somatosensory predictions that produce attenuation. This finding reaffirms that action might constitute the most efficient way to distinguish ourselves from others.

## Acknowledgements

Konstantina Kilteni was supported by the Marie Sk**ł**odowska-Curie Intra-European Individual Fellowship (#704438). The project was funded by the Swedish Research Council, the Go☐ran Gustafssons Stiftelse and the Torsten So☐derbergs Stiftelse. We thank Martti Mercurio for technical support.

## Author Contributions

Konstantina Kilteni and H. Henrik Ehrsson conceived and designed the experiment. Konstantina Kilteni and Patrick Engeler collected together the data of the experiment. Konstantina Kilteni conducted the statistical analysis. Konstantina Kilteni and H. Henrik Ehrsson wrote the manuscript, and Patrick Engeler read and approved the final version.

## Appendix

**Supplementary Figure S1.**
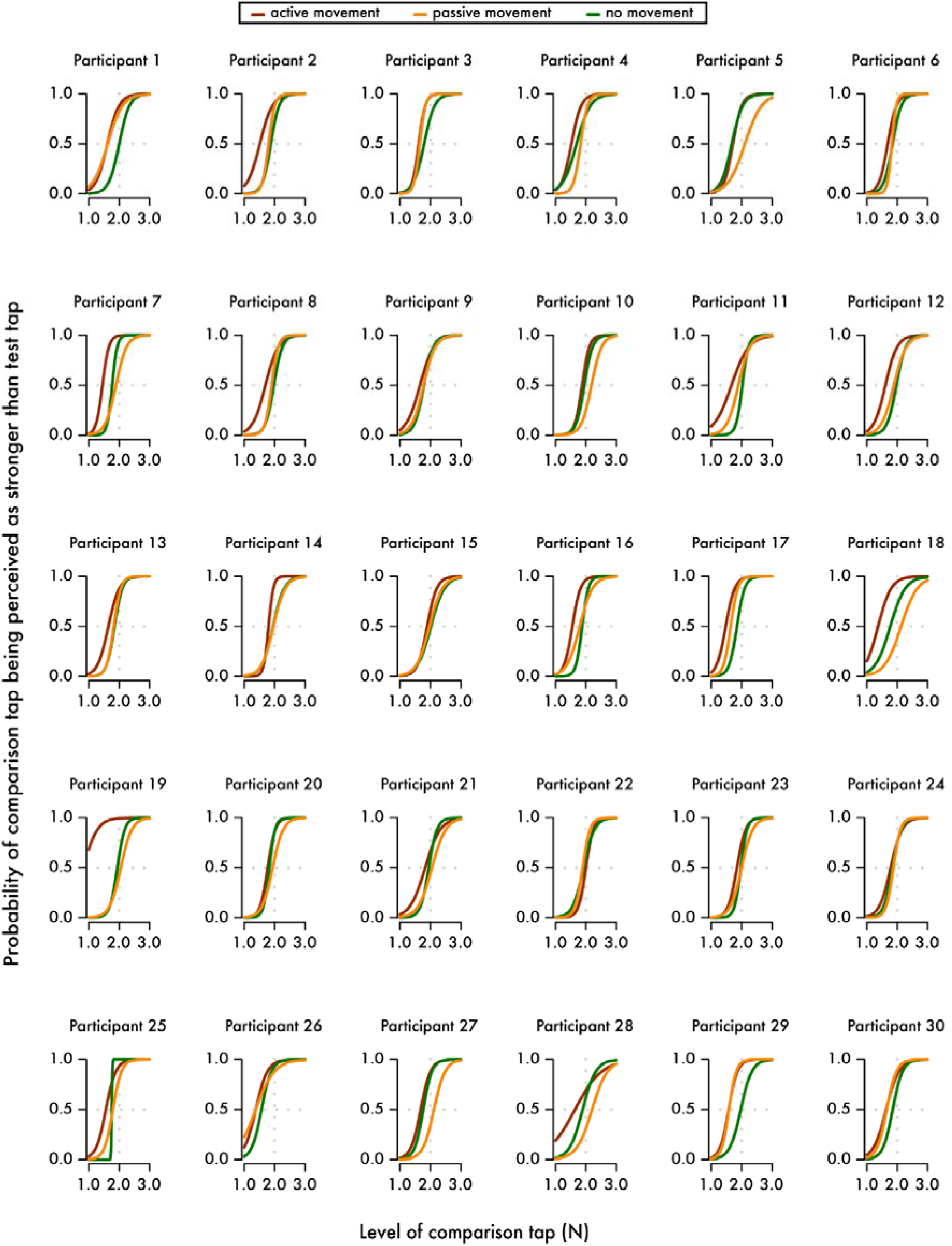
Individual fits per condition. Horizontal gray dotted lines indicate the 50% probability of the comparison tap being perceived as stronger than the test tap (PSE), while the vertical gray dotted lines indicate the true intensity of the test tap (2 N).

## References

Allen, M., Poggiali, D., Whitaker, K., Marshall, T.R. and Kievit, R.A. 2019. Raincloud plots: a multi-platform tool for robust data visualization. Wellcome Open Res., doi: 10.12688/wellcomeopenres.15191.1.

Bays, P.M., Flanagan, J.R. and Wolpert, D.M. 2006. Attenuation of self-generated tactile sensations is predictive, not postdictive. PLoS Biol., 4: 281–284.

Bays, P.M. and Wolpert, D.M. 2007. Computational principles of sensorimotor control that minimize uncertainty and variability. J. Physiol., 578: 387–396.

Bays, P.M. and Wolpert, D.M. 2008. Predictive attenuation in the perception of touch. In:. Sensorimotor Foundations of Higher Cognition (E. P. Haggard, Y. Rosetti, and M. Kawato, eds), pp. 339–358.

Bays, P.M., Wolpert, D.M. and Flanagan, J.R. 2005. Perception of the consequences of self-action is temporally tuned and event driven. Curr. Biol., 15: 1125–1128.

Blakemore, S.J., Frith, C.D. and Wolpert, D.M. 1999. Spatio-temporal prediction modulates the perception of self-produced stimuli. J. Cogn. Neurosci., 11: 551–559.

Blakemore, S.J., Wolpert, D.M. and Frith, C. 2000. Why can’t you tickle yourself? Neuroreport, 11: R11–R16.

Blakemore, S.J., Wolpert, D.M. and Frith, C.D. 1998. Central cancellation of self-produced tickle sensation. Nat. Neurosci., 1: 635–640.

Core Team, R. 2018. R: A language and environment for statistical computing.

Crapse, T.B. and Sommer, M.A. 2008. Corollary discharge across the animal kingdom.

Cullen, K.E. 2012. The vestibular system: Multimodal integration and encoding of self-motion for motor control.

Davidson, P.R. and Wolpert, D.M. 2005. Widespread access to predictive models in the motor system: a short review. J. Neural Eng., 2: S313–S319.

Diedrichsen, J., Verstynen, T., Hon, A., Lehman, S.L. and Ivry, R.B. 2003. Anticipatory adjustments in the unloading task: Is an efference copy necessary for learning? Exp. Brain Res., 148: 272–276.

Doemges, F. and Rack, P.M. 1992. Changes in the stretch reflex of the human first dorsal interosseous muscle during different tasks. J. Physiol., doi: 10.1113/jphysiol.1992.sp019018.

Franklin, D.W. and Wolpert, D.M. 2011. Computational mechanisms of sensorimotor control.

Friston, K. 2005. A theory of cortical responses. Philos. Trans. R. Soc. B Biol. Sci., doi: 10.1098/rstb.2005.1622.

Friston, K. 2009. The free-energy principle: a rough guide to the brain? Trends Cogn. Sci., 13: 293–301.

Grotheer, M. and Kovács, G. 2016. Can predictive coding explain repetition suppression? Cortex, 80: 113–124.

JASP and JASP Team. 2019. JASP.

Kawato, M. 1999. Internal models for motor control and trajectory planning. Curr Opin Neurobiol, 9: 718–727.

Kilteni, K., Andersson, B.J., Houborg, C. and Ehrsson, H.H. 2018. Motor imagery involves predicting the sensory consequences of the imagined movement. Nat. Commun., 9: 1617.

Kilteni, K. and Ehrsson, H. 2019. Functional connectivity between cerebellum and somatosensory areas reflects the attenuation of self-generated touch. Under Rev.

Kilteni, K. and Ehrsson, H.H. 2017a. Body ownership determines the attenuation of self-generated tactile sensations. Proc. Natl. Acad. Sci., 201703347.

Kilteni, K. and Ehrsson, H.H. 2017b. Sensorimotor predictions and tool use: Hand-held tools attenuate self-touch. Cognition, 165: 1–9.

Kilteni, K., Houborg, C. and Ehrsson, H.H. 2019. Rapid learning and unlearning of predicted sensory delays in self-generated touch. bioRxiv, 653923.

Oldfield, R.C. 1971. The assessment and analysis of handedness: the Edinburgh inventory. Neuropsychologia, 9: 97–113.

Palmer, C.E., Davare, M. and Kilner, J.M. 2016. Physiological and Perceptual Sensory Attenuation Have Different Underlying Neurophysiological Correlates. J. Neurosci., 36: 10803–10812.

Rao, R.P. and Ballard, D.H. 1999. Predictive coding in the visual cortex: a functional interpretation of some extra-classical receptive-field effects. Nat. Neurosci., 2: 79–87.

Shadmehr, R. and Krakauer, J.W. 2008. A computational neuroanatomy for motor control. Exp. Brain Res., 185: 359–381.

Shadmehr, R., Smith, M. a and Krakauer, J.W. 2010. Error correction, sensory prediction, and adaptation in motor control. Annu. Rev. Neurosci., 33: 89–108.

Shergill, S.S., Bays, P.M., Frith, C.D. and Wolpert, D.M. 2003. Two eyes for an eye: the neuroscience of force escalation. Science, 301: 187.

Shergill, S.S., Samson, G., Bays, P.M., Frith, C.D. and Wolpert, D.M. 2005. Evidence for sensory prediction deficits in schizophrenia. Am. J. Psychiatry, 162: 2384–2386.

Shergill, S.S., White, T.P., Joyce, D.W., Bays, P.M., Wolpert, D.M. and Frith, C.D. 2014. Functional magnetic resonance imaging of impaired sensory prediction in schizophrenia. JAMA psychiatry, 71: 28–35.

Therrien, A.S. and Bastian, A.J. 2018. The cerebellum as a movement sensor. Neurosci. Lett., 0–1.

Von Helmholtz, H. 1867. Handbuch der physiologischen Optik.

Walsh, L.D., Taylor, J.L. and Gandevia, S.C. 2011. Overestimation of force during matching of externally generated forces. J. Physiol., 589: 547–557.

Wolpe, N., Ingram, J.N., Tsvetanov, K.A., Geerligs, L., Kievit, R.A., Henson, R.N., Wolpert, D.M. and Rowe, J.B. 2016. Ageing increases reliance on sensorimotor prediction through structural and functional differences in frontostriatal circuits. Nat. Commun., 7.

Wolpert, D.M. and Flanagan, J.R. 2001. Motor prediction. Curr. Biol., 11: R729–R732.

Wolpert, D.M. and Ghahramani, Z. 2000. Computational principles of movement neuroscience. Nat. Neurosci., 3: 1212–1217.

Wolpert, D.M., Miall, R.C. and Kawato, M. 1998. Internal models in the cerebellum.

